# Patient reported distress can aid clinical decision making in idiopathic pulmonary fibrosis: analysis of the PROFILE cohort

**DOI:** 10.1101/460626

**Authors:** Iain Stewart, Tricia McKeever, Rebecca Braybrooke, Eunice Oballa, Juliet K Simpson, Toby M Maher, Richard P Marshall, Pauline T Lukey, William A Fahy, Gisli Jenkins, Gauri Saini

**Author notes:** Corresponding author: Iain Stewart, PhD; Division of Respiratory Medicine, University of Nottingham; B19, Clinical Sciences Building; Nottingham City Hospital; Nottingham; NG5 1PB.

## Abstract

Idiopathic pulmonary fibrosis is a progressive and fatal interstitial lung disease. We aimed to determine if patient response to a palliative assessment survey could predict disease progression or death.

We undertook a cross-sectional study in a UK clinical cohort of incident cases. Rasch-based methodology provided a disease distress value from an abridged 11 item model of the original 45 item survey. Distress values were compared with measures of lung function. Disease progression or mortality alone was predicted at twelve months from survey completion, with risk of death assessed at three, six and twelve months.

Disease distress values were negatively correlated with lung function (r=-0.275 percent predicted DLCO). Expected survey scores computed from distress values could distinguish disease progression, 8.8 (p=0.004), and people who died, 10.2 (p=0.002), from those who did not progress, 6.9. Actual survey scores predicted disease progression and mortality with an area under the curve of 0.60 and 0.64, respectively. Each point increment in actual score increased risk of twelve-month mortality by 10%, almost 43% of people scoring above 18 did not survive beyond 105 days.

We define a short questionnaire that can score disease distress and predict prognosis, assisting clinical decision making in progressive fibrosis.

## 1. Introduction

Idiopathic pulmonary fibrosis (IPF) is a progressive interstitial lung disease of unknown aetiology, causing scarring of the lung, shortness of breath, cough and reduction in lung function. A UK study demonstrated an increase in incidence of IPF by 35% between 2000 and 2008, with higher incidence in men and older age groups [1]. Incidence and mortality continues to rise and is also increasing globally [2]. IPF is fatal with no current cure, though disease-modifying treatments are being introduced. In many cases, earlier palliative intervention could reduce the burden on the individual, as recommended in cancer care [3, 4], yet the variable natural history of IPF makes it difficult to predict care needs.

The Sheffield Profile for Assessment and Referral to Care (SPARC) is a holistic needs tool, comprising 45 items across 9 domains, that can aid health professionals in identifying needs for palliative care [5]. Respondents find the questions easy to understand [6] and the tool has been adapted to international settings [7]. Such systematic assessment provides a useful indication of symptom distress [8]. Yet those who suffer chronic disease may not achieve the SPARC criteria for immediate clinical assessment, defined as a score of 3 in a single question, as habituation or affective comorbidity can affect symptom perception [5, 9, 10]. Regular completion of 45 items can be burdensome for those who suffer most [8], whilst self-assessment may result in interpatient differences despite the same underlying distress.

Rasch-based methodology generates a scaled value for a set of responses as an interaction between question difficulty and the individuals’ likelihood of scoring. Initially used to standardise scholastic tests, extension of the Item Response Theory (IRT) class of models to allow multiple choice or graded options has found increased recognition in developing healthcare metrics and patient-reported outcomes [11-15]. Further advantages of the method enables question refinement, banking and assessment of the impact of demographics on item response [16-18].

We aimed to determine whether the SPARC could be used with IRT methodology to generate a tool that could appropriately distinguish disease progression in a UK clinical cohort of IPF patients [19, 20], helping to predict short-term prognosis.

## 2. Methods

The Prospective Observation of Fibrosis in Lung Clinical Endpoints (PROFILE) Central England (NCT01134822) study is a longitudinal observational clinical trial that has been described previously [19, 20]. Age was grouped at 65 or under, 66-79, 80 or over; comorbidities were grouped as none, 1-2, 3 or more. People were asked to complete the SPARC questionnaire, comprised of dichotomous items in two domains, as well as polytomous items in six domains (Supplementary Doc 1). 243 people from the PROFILE Central England study completed the SPARC and were subsequently included in the analysis.

All analyses were performed in Stata (SE15.1). Lung function measures recorded at the initial SPARC assessment (±30 days) were used as baseline, with an outcome of disease progression within one year of SPARC completion defined as 10% relative decline in FVC or death. Percent predicted DLCO (%DLCO) and FVC (%FVC) was calculated using the suite of Global Lung Function Initiative (GLI) tools available from the European Respiratory Society [21]. Where %FVC (n=21) or %DLCO (n=75) could not be confirmed due to missing data, people were excluded from the relevant statistical analyses.

A two-parameter graded response model was ultimately constructed with items showing good discrimination and model assumptions were verified [11, 22]. Parameters of discrimination and difficulty were assigned to each item (survey question) according to how well it could differentiate people across the scale of the underlying distress trait (theta), as well as the probability of a particular answer. Detailed information on the construct of the IRT model is provided as supplementary information; a final model of 11 items was built, where larger theta values indicate more distress. (Supplementary Doc 2; Supplementary Table 1; Supplementary Figure SF1).

Pearson’s Correlation determined whether distress (theta) values correlated with %DLCO or %FVC measures. %DLCO was log transformed to meet normality assumptions. IRT test characteristic curves estimated expected questionnaire scores according to distress values calculated from the concise 11 item model, hereafter termed IPF Prognostic Assessment and Referral to Care (IPARC). One-way ANOVA assessed differences in mean distress between categories of %DLCO (<40%, 40-60%, >60%). Two-way t-test assessed mean distress between those with disease progression and those without, whilst one-way ANOVA additionally assessed categories of disease progression (no disease progression, lung function decline only, death only). Tukey post-hoc analysis between categories was applied.

We calculated the area under the Receiver Operator Characteristic (ROC) curve for the ability of the cumulative IPARC Score to predict disease progression within 12 months, as well as overall mortality compared with %FVC or %DLCO. The sensitivity and the specificity was compared using the chi-squared test [23].

Kaplan-Meier curves plotted time to disease progression, or overall mortality, against days since completing the questionnaire according to categories of IPARC Score. Cox regression estimated the hazard ratio (HR) of disease progression, as well as death in 365 days (12 months), 180 days (6 months) and 90 days (3 months) according to increment or categories of IPARC Scores; lowest scorers as reference. The risk in comparison groups is contingent upon the proportion of the reference group that fails during the specified timescale. Analyses were initially univariate, and then adjusted for age, comorbidity, gender, and a significant interaction between age and comorbidity. Proportional hazard assumptions were checked using Schoenfeld residuals.

## 3. Results

### Demographics

From 243 people within the cohort, 103 (42.4%) had evidence of disease progression within one year, whilst 140 (57.6%) did not (Table 1). Of those with disease progression, 49 died within one year (47.6%). Within the disease progression subgroup, 80.6% were male but no significant gender interaction was observed, similarly comorbidity count was not associated with progression. Those with disease progression had greater representation by ages 80 or over. We identified no significant relationships between communication and progression status, whilst we observe lower proportions with disease progression wanting information about personal finances. No relationship reached significance following Bonferroni correction.

**Table 1:**
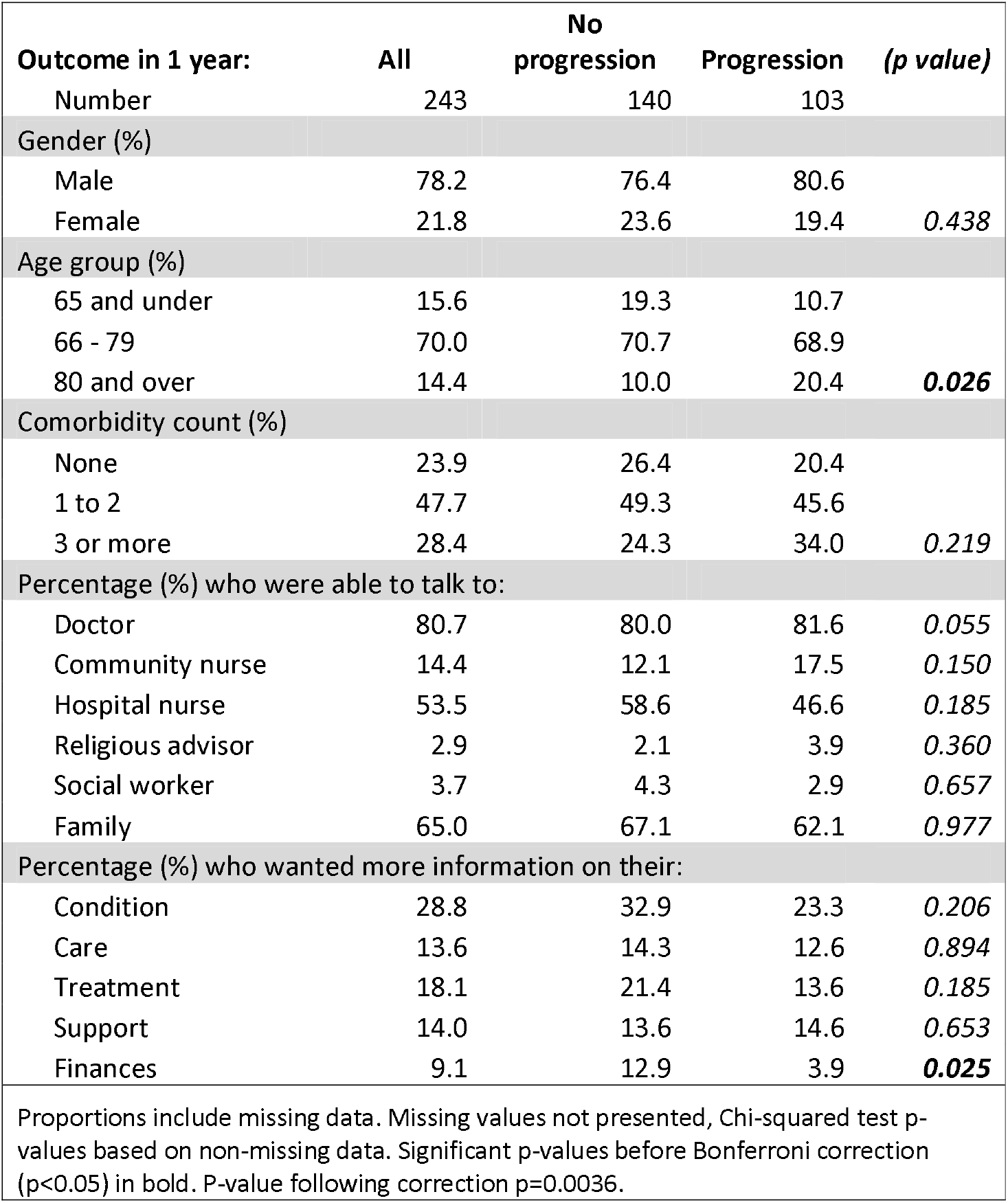
Cohort demographics

### Disease distress is associated with lung function and disease progression

A mild negative correlation was observed between distress values and baseline %FVC or %DLCO, indicating that higher distress correlated with worse lung function performance (Figure 1). The Pearson correlation coefficient for distress and %FVC was -0.267 (p=0.0001), whilst for distress and %DLCO it was -0.275 (p=0.0003).

**Figure 1.**
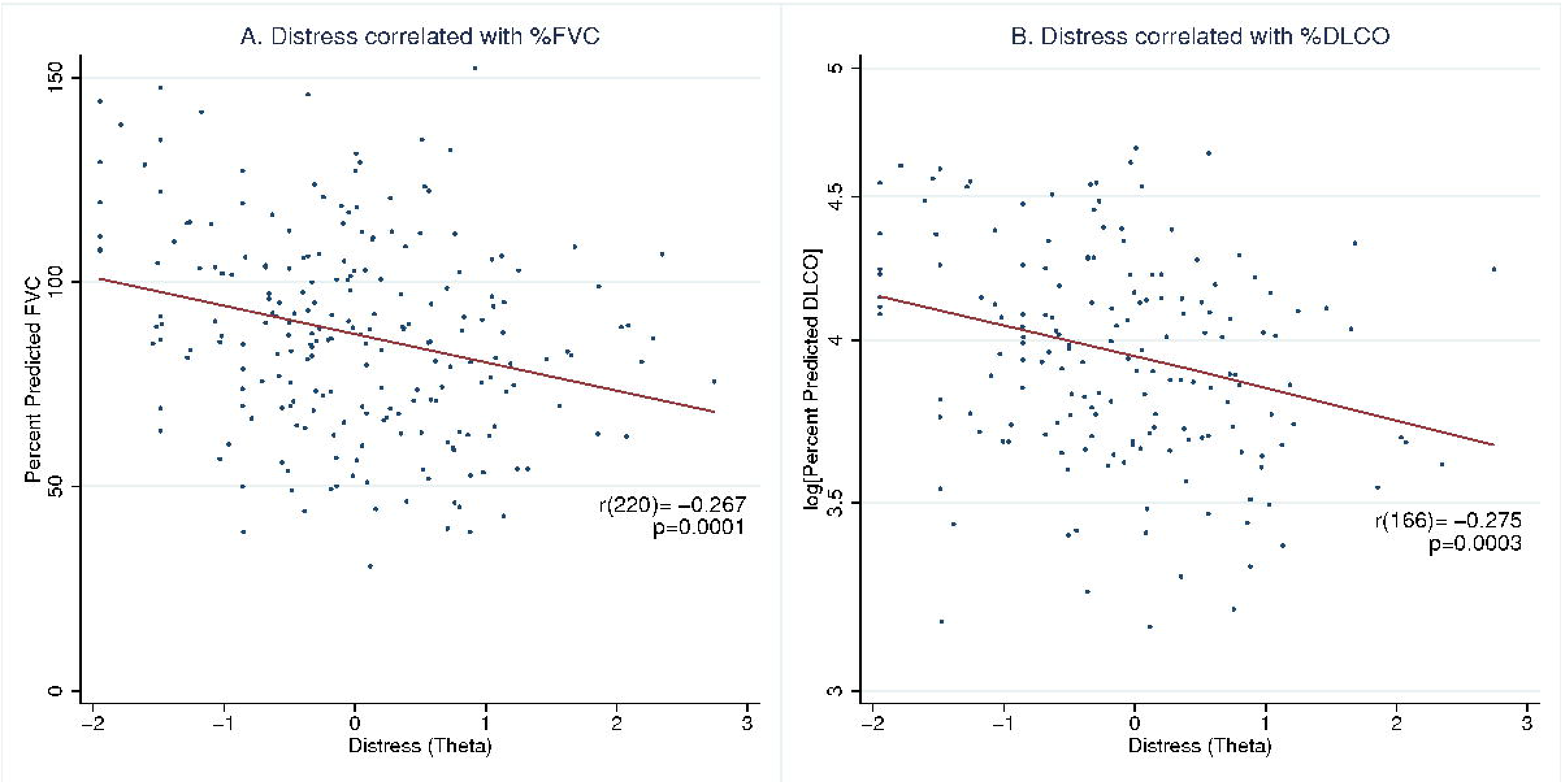
Negative correlation of distress with percent predicted lung function. A) Plotted %FVC against distress value generated from IPARC Score model resulted in a Pearson correlation coefficient (r) of -0.267 (p=0.0001) from 222 people (220 degrees freedom); %FVC explains 7.1% of variation in disease distress. B) Plotted Log transformed %DLCO against distress value resulted in r=-0.275 (p=0.0003) from 168 people (166 degrees freedom); %DLCO explains 7.6% of variation in disease distress.

The mean distress value according to severity of %DLCO was calculated and the IRT test characteristic curve estimated the expected score from the continuum of the distress values, calculated from the IPARC Score model. Mean distress values were significantly different according to severity of %DLCO (p=0.006). People with %DLCO <40% were significantly more distressed than those with a %DLCO >60% (p=0.004), leading to expected scores of 9.1 (θ=0.22) and 5.6 (θ=-O.38), respectively (Figure 2 A).

**Figure 2.**
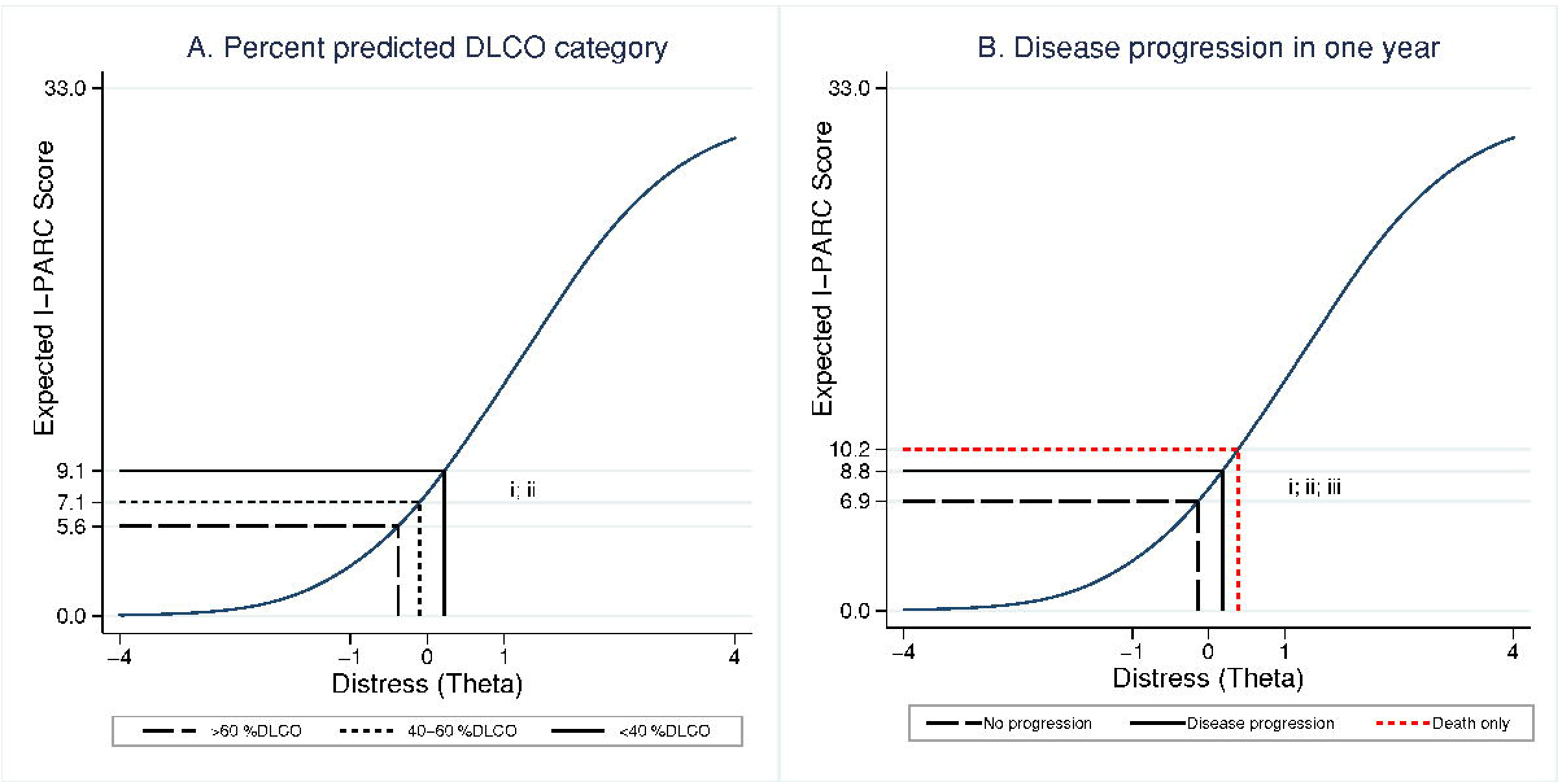
Expected IPARC Scores from mean distress in lung function categories. Item response theory test characteristic curve from 11 items in final model plots distress (theta) values against expected score. Scale of theta has a mean of 0 and the arbitrary range (4) represents incremental standard deviations. A) Mean theta values plotted for %DLCO category: >60% (long dash), 40-60% (short dash), <40% (solid); i. one-way ANOVA p=0.006; ii. Post-hoc Tukey analysis p=0.004 <40 %DLCO compared with >60. B) Mean theta values plotted for disease progression: no progression (long dash), disease progression (solid); i. Two way t-test p=0.0085. Death only (short dash); One-way ANOVA death, decline only, no decline p=0.003; iii. Post-hoc Tukey analysis p=0.002 death compared with no decline.

Mean distress values were also significantly different according to disease progression (p=0.0085), where those who progressed had an expected score of 8.8 (θ=0.18) compared to an expected score of 6.9 for those who did not progress (0=-O.14) (Figure 2 B-i). Plotting mean distress of patients that died separately provided an expected score of 10.2 (θ=0.39); significance was identified across categories of disease progression (p=0.003), largely driven by the difference between those who died and those who did not progress (p=0.002) (Figure 2 Bii-iii).

Lung function recordings require a suitable fitness to provide acceptable and repeatable measurements, introducing selection bias if individuals struggle to perform them. To understand the influence of missing data we evaluated distress according to completion of lung function (see Supplementary Table 2). Patients with missing %FVC data were more likely to have died (61.9%) than patients with complete %FVC data (16.2%; p<0.0001). However, there was no difference in mean distress between people with missing %FVC compared with complete data (p=0.384, compare scores 8.8 versus 7.6). Similarly, patients with missing %DLCO data were more likely to have died (34.7% versus 13.7%; p <0.0001), although those missing the data were also more distressed (p=0.002; compare scores 9.5 versus 7.0). In total, 75 people (31%) were missing DLCO measures; analyses based on non-missing DLCO recordings will underestimate actual effect sizes as they exclude a large sample of distressed patients.

### IPARC Score can predict disease progression and mortality

Area under the curve (AUC) of the ROC curve initially assessed the ability of the IPARC Score to predict disease progression in the complete dataset (Figure 3), resulting in an AUC of 0.60 (95%CI: 0.53-0.68). ROC subsequently assessed the ability of the IPARC Score to predict death compared with lung function recordings in 167 people with complete data. We identified %DLCO as having the largest AUC of 0.83 (95%CI: 0.75-91), whilst %FVC had an AUC of 0.66 (95%CI: 0.56-0.76). The cumulative IPARC Score provided an AUC of 0.64 (95%CI: 0.52-0.77); a minimum score of 9 resulted in 59.2% sensitivity and 62.9% specificity. No statistical difference was observed between the AUC of IPARC Score and %FVC (p=0.8), although %DLCO was significantly better at predicting mortality (p=0.013). Ability of IPARC Score to predict mortality was also assessed independently from lung function recordings in the complete dataset (see Supplementary Figure SF2), resulting in a greater AUC of 0.66 (95%CI: 0.57-0.74).

**Figure 3.**
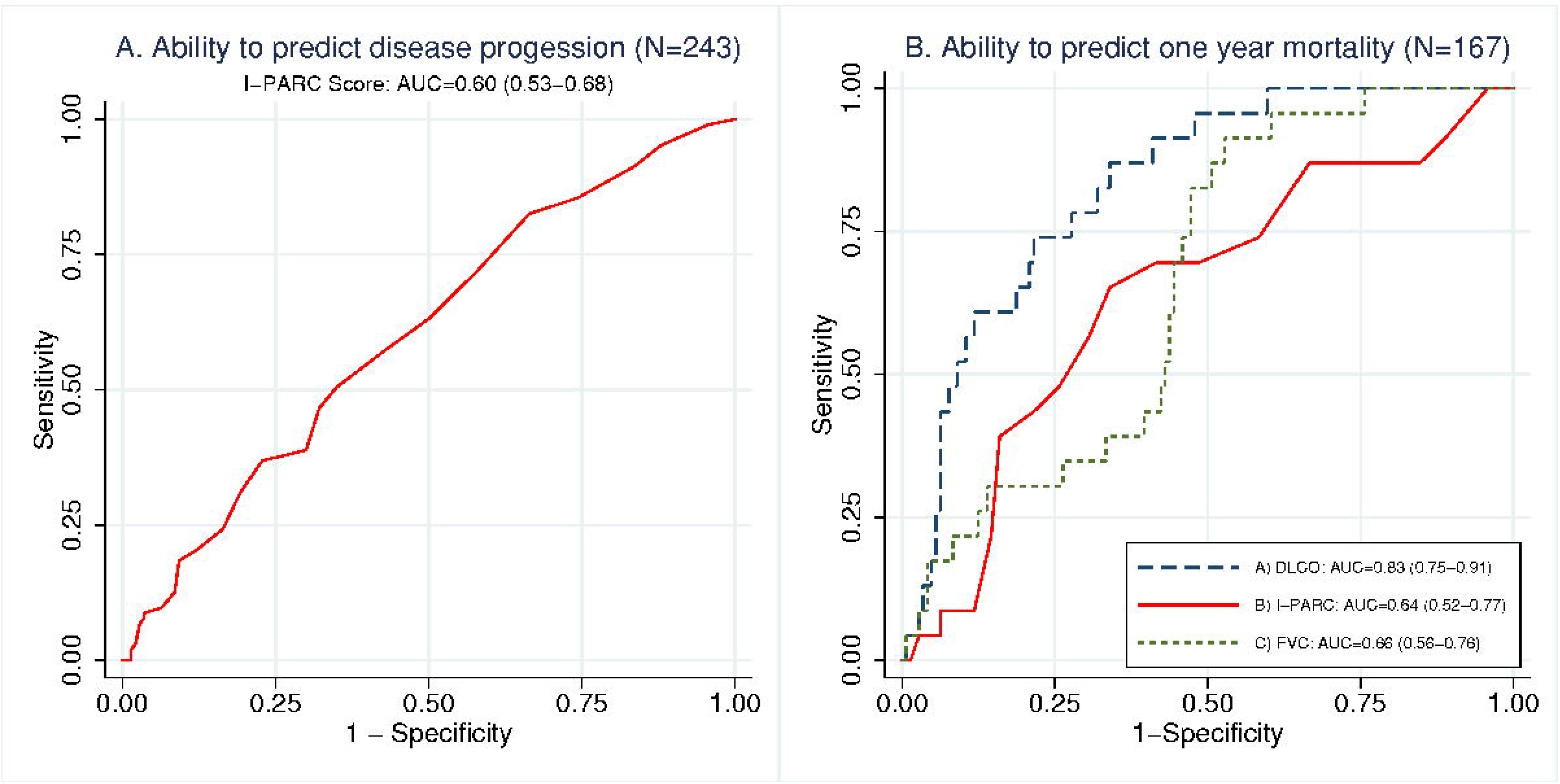
Receiver-Operator Curve (ROC) assesses sensitivity and specificity of measures in predicting progression. A) Area under curve (AUC) for IPARC Score in predicting disease progression = 0.60 (95% confidence interval 0.53-0.68) B. AUC for IPARC Score (solid line) in predicting one year mortality =0.64 (0.52-0.77); %DLCO (long dash) AUC=0.83 (0.75-0.91); %FVC (short dash) AUC=0.66 (0.56-0.76). %DLCO AUC greater than IPARC AUC, p=0.013.

Kaplan Meier curves of time to event were plotted separately for disease progression and death according to observed IPARC Score (Figure 4), categorised using expected score from mean distress where % DLCO <40% (score of 9). A large proportion of people scoring over 18 from the 11 IPARC items had rapid disease progression, 51.5% of people scoring at least 9 progressed within one year. Fewer than 75% of people who scored between 9 and 18 survived 365 days (Figure 4 B). A total of 101 people scored 9 or more and 29 died within 365 days, providing a positive predictive value of 28.7%; this is compared with 20 people from the 142 scoring 8 or less, 14.1%. Scoring over 18 increased the positive predictive value to 42.9% from a total of 14 people, all deaths in this group occurred within 105 days.

**Figure 4.**
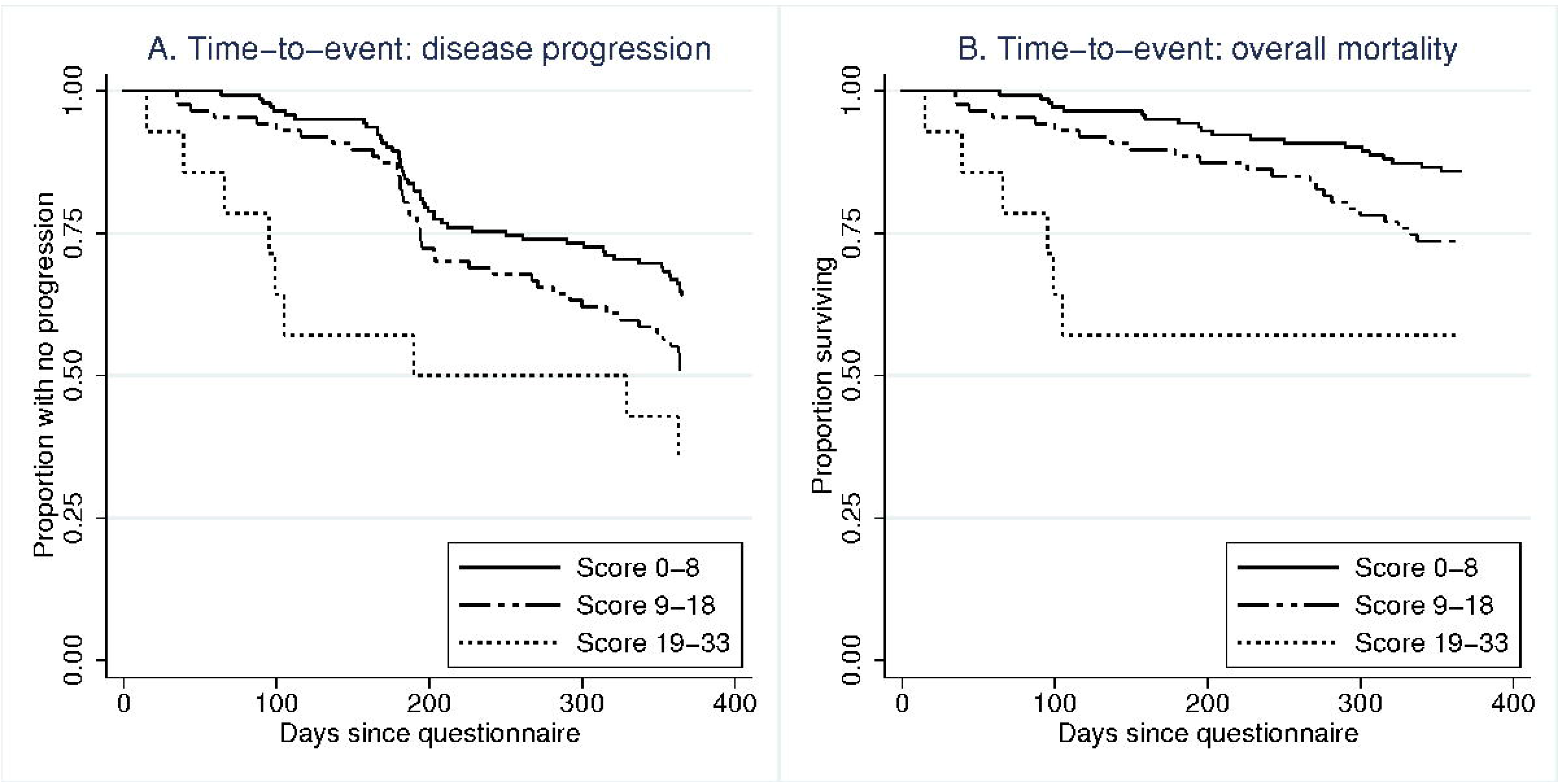
Kaplan Meier analyses of time to event. A) Kaplan Meier Curves plot proportion of people with no disease progression against time since completing questionnaire: IPARC Score low (0-8, solid line), intermediate (8-18, dashed), high (9-33, dotted). B) Kaplan Meier Curves plot proportion of people surviving against time since completing questionnaire: IPARC Score low (0-8, solid line), intermediate (8-18, dashed), high (9-33, dotted).

Cox regression estimates of the risk of disease progression and risk of death in one year for each unit increment of the IPARC Score are reported in Table 2. For every point increase, the risk of disease progression increased by 5% (HR 1.05 95%CI 1.02-1.09) whilst risk of death increased by 10% (HR 1.10 95%CI 1.05-1.15). Disease progression was almost 60% more likely for people scoring 9-18 (HR 1.59 95%CI 1.04-2.44), compared to those scoring less, whilst those scoring highest were at three times the risk (HR 3.19 95%CI 1.53-6.64). People scoring between 9 and 18 were more than twice as likely to die within 365 days as those scoring less than 9 (HR 2.43 95%CI 1.30-4.55). Estimates for those scoring over 18 did not meet proportional hazard assumptions at 365 days, although their risk of death in 180 days was 12 fold that of those scoring less than 9 (HR 12.32 95%CI 3.84-39.55), whilst the risk for those scoring between 9 and 18 more than doubled (HR 2.82 95%CI 1.03-7.68). Similar results were estimated at 90 days for those scoring highest (HR 19.47 95%CI 1.98-191.13), though scoring between 9 and 18 was not statistically different from scoring less (HR 7.69 95%CI 0.89-66.28).

**Table 2.** Cox regression risk estimates of disease progression and overall mortality by IPARC Score.

|  | Number | % fail | HR | 95%CI | HR* | 95%CI* |
| --- | --- | --- | --- | --- | --- | --- |
| **365 day disease progression risk** |  |  |  |  |  |  |
| *IPARC Score* | 243 | 42.39% | 1.05 | (1.01-1.08) | 1.05 | (1.02-1.09) |
| *IPARC category* |  |  |  |  |  |  |
| Score 0-8 | 142 | 35.92% | 1 |  | 1 |  |
| Score 9-18 | 87 | 49.43% | 1.50 | (1.00-2.26) | 1.59 | (1.04-2.44) |
| Score 19-33 | 14 | 64.29% | 2.69 | (1.32-5.47) | 3.19 | (1.53-6.64) |
| **365 day mortality risk** |  |  |  |  |  |  |
| *IPARC Score* | 243 | 20.16% | 1.08 | (1.03-1.12) | 1.10 | (1.05-1.15) |
| *IPARC category* |  |  |  |  |  |  |
| Score 0-8 | 142 | 14.08% | 1 |  | 1 |  |
| Score 9-18 | 87 | 26.44% | 2.02 | (1.11-3.68) | 2.43 | (1.30-4.55) |
| Score 19-33 | 14 | 42.86% | 4.44 | (1.78-11.08) | †5.74 | (2.20-14.97) |
| **180 day mortality risk** |  |  |  |  |  |  |
| *IPARC category* |  |  |  |  |  |  |
| Score 0-8 | 142 | 4.93% | 1 |  | 1 |  |
| Score 9-18 | 87 | 11.49% | 2.43 | (0.92-6.37) | 2.82 | (1.03-7.68) |
| Score 19-33 | 14 | 42.86% | 11.69 | (3.92-34.86) | 12.32 | (3.84-39.55) |

| 90 day mortality risk |  |  |  |  |  |  |
| --- | --- | --- | --- | --- | --- | --- |
| *IPARC category* |  |  |  |  |  |  |
| Score 0-8 | 142 | 0.70% | 1 |  | 1 |  |
| Score 9-18 | 87 | 5.75% | 8.37 | (0.98-71.61) | 7.69 | (0.89-66.28) |
| Score 19-33 | 14 | 21.43% | 34.25 | (3.56-329.37) | 19.47 | (1.98-191.13) |
| *adjusted for age group, comorbidity count, gender, age comorbidity interaction.<br>Proportional hazard assumptions met unless stated (†). HR: hazard ratio |  |  |  |  |  |  |

A sub-analysis was performed on a small sample who had an IPARC score before and after pirfenidone treatment to preliminarily assess therapy modification of the IPARC score (Supplementary Figure SF3). Overall, in 12 patients with matched scores taken at a year interval, there was a non-significant trend towards increased distress following therapy with pirfenidone. The change in level of distress with treatment was driven by the IPARC category prior to therapy; patients scoring under 9 (mild distress) demonstrated more distress following therapy (p=0.0086), whilst those scoring 9 or over (high level of distress) had a more varied response including some reduction in IPARC score.

## 4. Discussion

The SPARC questionnaire is recognised within the UK National Health Service as a way to help address and improve end-of-life care management [5, 6, 9, 24]. We have utilised a unique clinical cohort of patients with IPF [19], determining that items specified within the IPARC list can identify patients at high risk of death within 3-6 months, providing opportunity for earlier supportive care and improving patient outcomes (Box 1).

#### Box 1: Scoring of items in final model

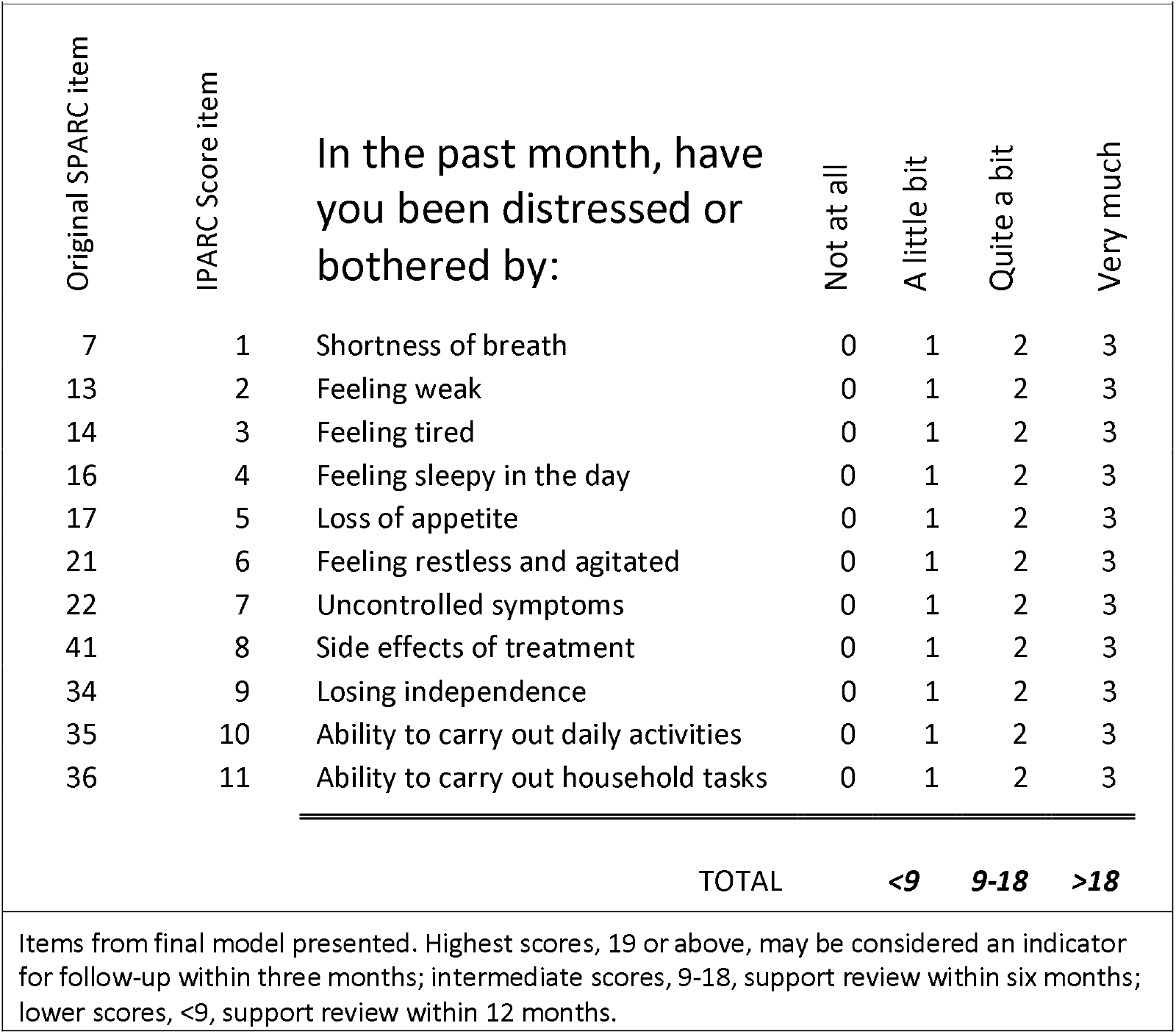

We use Item Response Theory as a novel application of the methodology to assess patient reported outcome measures [11], identifying a concise list of 11 items from an original 45 questions that capture the majority of information distinguishing people with high distress. Disease distress correlated with percent predicted lung function and could characterise disease progression. Furthermore, scores could predict mortality with those scoring highest also at greatest risk of death. The associations of disease distress with lung function, disease progression and mortality support the IPARC Score as a valid tool to inform prognoses.

Previous studies using alternate patient reported outcomes to measure quality of life in IPF have similarly noted weak to moderate correlations with lung function measures, including the MRC dyspnoea scale, ATAQ-IPF and CAT [25-27]. Imperfect correlations indicate that lung function alone cannot account for all differences in patient reported outcomes, in combination they offer tremendous clinical value [28]. However, the MRC dyspnoea scale is one dimensional, ATAQ-IPF remains extensive at 75 items and CAT was developed for COPD, which at 8 items is convenient but may not have the sensitivity to capture IPF traits [29]. Whilst the original SPARC is a broad palliative assessment survey, answers provided by an IPF population specify appropriate items in the IPARC tool. The unabbreviated name, IPF Prognostic Assessment and Referral to Care, reflects the purpose and cohort it was developed with, as well as the tool it originated from.

### Strengths and limitations of Item Response Theory application

IRT is an increasingly applied methodology that lends itself to parametric tests, providing an evidence base for reusing questions that can be used to distinguish traits of interest [16, 18, 30]. This study supports the use of IRT methodology in optimising questions within healthcare surveys to predict patient outcomes. Questions that did not distinguish levels of distress, such as those which scored low for the vast majority (e.g. Q33: Religious or spiritual needs not being met), or those that can show clustering of responses (e.g. Q30: Thoughts about ending it all; Q32: Worrying thoughts about death or dying) can be dropped in order to retain items offering the most discriminatory value.

IRT methodology can also assess the way in which an item may be answered differently according to demographic traits that are unrelated to the latent trait of distress, termed differential item functioning (DIF)[16, 17]. We identified no DIF in the final model according to gender, indicating men and women with IPF report distress similarly. Of the 243 individuals sampled in this study two were of non-white origin, and thus we were not able to assess the influence of ethnicity on patient responses. Further study in more ethnically diverse populations is warranted to determine whether IPARC retains prognostic value. DIF by demographics of age category and comorbidity count was identified for item 21 (Feeling restless and agitated), however the analyses indicate an underlying relationship between these demographics and disease distress, so are not defined as DIF. The original SPARC questionnaire collects distress information from an extensive list of possible issues, with a response of ‘very much’ (score of 3) in any of the 41 scaled items acting as a flag that the individual would benefit from an immediate palliative care assessment. Within clinical settings, the SPARC questionnaire could be considered too sensitive, or excessive depending on health status [6, 9], whilst underreporting of distress is an issue in progressive disease [10, 31]. Use of the cumulative score over 11 items provides greater opportunity to capture distress when underreported, whilst shorter health questionnaires can provide similar insight to lengthier ones [32].

The mean distress for those with lowest %DLCO equated to a score of 9.1, with a score of 9 being subsequently used as a threshold point for grouping people who may undergo disease progression within one year. Lung function measures are a valuable tool in defining disease progression, although limited by the scheduling of measurement recordings that may not reflect the actual timing of progression. Those who progressed had an average score of 8.8, which supports an IPARC Score of 9 as a suitable benchmark, particularly as those who died had an average score of 10.2.

Other self-reported surveys exist to measure health status in those with progressive lung disease, including King’s-Brief Interstitial Lung Disease (K-BILD) questionnaire[15]. The authors of K-BILD used patient interviews to define a series of pertinent questions and Rasch analysis to refine the item list to 15 with 4 domains and a 7-point Likert scale. We use similar methodology to refine the list of predetermined SPARC items, as answered by a sample of 243 IPF patients, to a set of 11 within a single domain and a 4-point Likert scale. Whilst K-BILD offers an empowering way to self-monitor respiratory health, IPARC should be completed within clinical settings where rapid disease progression is a factor.

### IPARC Score offers prognosis estimates for clinical decisions

ROC analysis confirmed that %DLCO was the best measure for predicting one year mortality in this cohort [33]. IPARC Score appeared to perform slightly better at identifying true positives compared with %FVC when the accepted proportion of false positives was restricted to 50%. Lung function measurements can be challenging for patients with severe disease to provide, and this appears particularly apparent in people missing DLCO who were significantly more distressed than those without missing DLCO values. These data illustrate the considerable survival bias associated with lung function data in studies of patients with progressive lung disease. As a result, these analyses may underestimate IPARC performance relative to lung function measures, yet demonstrate the value of patient-reported measures.

Therapy modified quality of life measures can inform whether an intervention is necessary or successful [34]. We undertook a preliminary analysis on a very small subset of patients, which indicated that treatment with pirfenidone in patients with mild levels of distress may be associated with increased levels of distress following anti-fibrotic therapy. This is consistent with the known adverse effect profile of pirfenidone though we do not account for clinical presentation or disease progression on therapy. We recommend further trials measure distress modification following therapeutic intervention to assess whether IPARC can be used to define individuals for whom a particular therapy may be heightening distress.

Survival curves indicated that whilst the majority of the cohort survived one year after completing their SPARC questionnaire, those with the highest IPARC scores had poorer life expectancy than those scoring lowest. The positive predictive value of mortality in one year was 29% for those scoring above the threshold of 9, and increased to 43% for those scoring 19 or more. The findings provide evidence that patient reported distress can predict early mortality with similar accuracy to predictions made with FVC lung function recordings.

Adjusted survival analyses further show that risk of one-year mortality increased by 10% with each incremental point, whilst risk of progression increased by 5%. Intermediate scoring of 9 to 18 heightened the likelihood of progression and more than doubled the risk of death, compared to those scoring lower. It can be observed that a large proportion of the highest scorers did not survive beyond 105 days, although these estimates are based on low numbers. This relatively large sample of people with confirmed IPF supports the benefit of patient reported distress in predicting disease progression and death.

Given that it is challenging to perform lung function in patients with progressive disease it is important to develop clinical markers of prognosis as well as assessing health status. The IPARC Score requires no specialist equipment or training to complete or calculate, takes consideration of the individual’s concerns, can be undertaken at any time without a requirement for repeated measures, and can aid decisions to review patients earlier. Combined with its associations with lung function and mortality, this patient-reported outcome is a potential component of composite endpoints in clinical trials, although further validation and studies on treatment sensitivity are essential [35]. The predictive capacity could not exclude all false positives, yet we recognise that early integration of palliative care can improve outcomes in other progressive diseases [3, 4, 36]. Future studies may include IPARC Score as part of a composite scoring system for accurately predicting IPF prognosis, adding value in treatment recommendations [33]. We recommend utilising it in combination with available lung function recordings when making clinical decisions (see Box 1).

The simple practicality of the tool allows an assessment of factors impacting quality of life to identify appropriate clinical management strategies such as supplemental oxygen for those reporting high distress with shortness of breath, or domiciliary support and assistive living devices to reduce distress from losing independence. Where best supportive care is indicated through high score and clinical presentation, the tool encourages a focus on the most distressing features for the individual [28]. We welcome further study to validate the IPARC tool in separate cohorts of patients with progressive lung disease, including IPF.

## Conclusion

The IPARC Score, developed using the SPARC holistic tool, offers an encouraging method to assess prognosis and recognise palliative care needs for those with progressive disease. In producing a standardised model based on the latent trait of distress for people with IPF, we generated an abridged list of items that could distinguish those who died within one year, with higher scores having worse prognosis. This brief and simple tool offers utility in the clinical care of patients with progressive lung fibrosis, whilst future study should address its validity following anti-fibrotic therapy and in other progressive lung diseases.

## Supporting information

