## Supplementary material for "Patient reported distress can aid clinical decision making in idiopathic pulmonary fibrosis: analysis of the PROFILE cohort"

University of Nottingham & Nottingham
Respiratory Biomedical Research Unit

### SPARC *

We would like to know a bit more about you and your concerns.

Please fill in this questionnaire (with help from a relative or carer if needed) and return it to one of our team.

There are no "right" or "wrong" answers. If you are unsure of a question, please leave it blank.

THANK YOU

* Sheffield Profile for Assessment and Referral to Care

Your initials:………………. Date completed:....../……./…….

#### COMMUNICATION AND INFORMATION ISSUES

**1. Have you been able to talk to any of the following people about your condition?**

| | Yes | No |
|---|---|---|
| a. Your doctor | ☐ | ☐ |
| b. Community nurse | ☐ | ☐ |
| c. Hospital nurse | ☐ | ☐ |
| d. Religious advisor | ☐ | ☐ |
| e. Social worker | ☐ | ☐ |
| f. Family | ☐ | ☐ |
| g. Other people (please state): ______________________ | | |

#### PHYSICAL SYMPTOMS

Please circle one answer per line

| ***In the past month, have you been distressed or bothered by:*** | **Not at all** | **A little bit** | **Quite a bit** | **Very much** |
|---|---|---|---|---|
| **2.** Pain? | 0 | 1 | 2 | 3 |
| **3.** Loss of memory? | 0 | 1 | 2 | 3 |
| **4.** Headache? | 0 | 1 | 2 | 3 |
| **5.** Dry mouth? | 0 | 1 | 2 | 3 |
| **6.** Sore mouth? | 0 | 1 | 2 | 3 |
| **7.** Shortness of breath? | 0 | 1 | 2 | 3 |
| **8.** Cough? | 0 | 1 | 2 | 3 |
| **9.** Feeling sick (nausea)? | 0 | 1 | 2 | 3 |
| **10.** Being sick (vomiting)? | 0 | 1 | 2 | 3 |
| **11.** Bowel problems (e.g. constipation, diarrhoea, incontinence)? | 0 | 1 | 2 | 3 |
| **12.** Bladder problems (urinary incontinence)? | 0 | 1 | 2 | 3 |
| **13.** Feeling weak? | 0 | 1 | 2 | 3 |
| **14.** Feeling tired? | 0 | 1 | 2 | 3 |
| **15.** Problems sleeping at night? | 0 | 1 | 2 | 3 |
| **16.** Feeling sleepy during the day? | 0 | 1 | 2 | 3 |

| PHYSICAL SYMPTOMS continued | | Not at all | A little bit | Quite a bit | Very much |
|---|---|---|---|---|---|
| **17.** | Loss of appetite? | 0 | 1 | 2 | 3 |
| **18.** | Changes in your weight? | 0 | 1 | 2 | 3 |
| **19.** | Problems with swallowing? | 0 | 1 | 2 | 3 |
| **20.** | Being concerned about changes in your appearance? | 0 | 1 | 2 | 3 |
| **21.** | Feeling restless and agitated? | 0 | 1 | 2 | 3 |
| **22.** | Feeling that your symptoms are not controlled? | 0 | 1 | 2 | 3 |

**PSYCHOLOGICAL ISSUES**

Please circle one answer per line

| ***In the past month, have you been distressed or bothered by:*** | | Not at all | A little bit | Quite a bit | Very much |
|---|---|---|---|---|---|
| **23.** | Feeling anxious? | 0 | 1 | 2 | 3 |
| **24.** | Feeling as if you are in a low mood? | 0 | 1 | 2 | 3 |
| **25.** | Feeling confused? | 0 | 1 | 2 | 3 |
| **26.** | Feeling as if you are unable to concentrate? | 0 | 1 | 2 | 3 |
| **27.** | Feeling lonely? | 0 | 1 | 2 | 3 |
| **28.** | Feeling that everything is an effort? | 0 | 1 | 2 | 3 |
| **29.** | Feeling that life is not worth living? | 0 | 1 | 2 | 3 |
| **30.** | Thoughts about ending it all? | 0 | 1 | 2 | 3 |
| **31.** | The effect of your condition on your sexual life? | 0 | 1 | 2 | 3 |

**RELIGIOUS AND SPIRITUAL ISSUES**

Please circle one answer per line

| ***In the past month, have you been distressed or bothered by:*** | | Not at all | A little bit | Quite a bit | Very much |
|---|---|---|---|---|---|
| **32.** | Worrying thoughts about death or dying? | 0 | 1 | 2 | 3 |
| **33.** | Religious or spiritual needs not being met? | 0 | 1 | 2 | 3 |

#### INDEPENDENCE AND ACTIVITY

Please circle one answer per line

| *In the past month, have you been distressed or bothered by:* | | Not at all | A little bit | Quite a bit | Very much |
|---|---|---|---|---|---|
| **34.** | Losing your independence? | 0 | 1 | 2 | 3 |
| **35.** | Changes in your ability to carry out your usual daily activities such as washing, bathing or going to the toilet? | 0 | 1 | 2 | 3 |
| **36.** | Changes in your ability to carry out your usual household tasks such as cooking for yourself or cleaning the house? | 0 | 1 | 2 | 3 |

#### FAMILY AND SOCIAL ISSUES

Please circle one answer per line

| *In the past month, have you been distressed or bothered by:* | | Not at all | A little bit | Quite a bit | Very much |
|---|---|---|---|---|---|
| **37.** | Feeling that people do not understand what you want? | 0 | 1 | 2 | 3 |
| **38.** | Worrying about the effect that your illness is having on your family or other people? | 0 | 1 | 2 | 3 |
| **39.** | Lack of support from your family or other people? | 0 | 1 | 2 | 3 |
| **40.** | Needing more help than your family or other people could give? | 0 | 1 | 2 | 3 |

#### TREATMENT ISSUES

Please circle one answer per line

| *In the past month, have you been distressed or bothered by:* | | Not at all | A little bit | Quite a bit | Very much |
|---|---|---|---|---|---|
| **41.** | Side effects from your treatment? | 0 | 1 | 2 | 3 |
| **42.** | Worrying about long term effects of your treatment? | 0 | 1 | 2 | 3 |

#### PERSONAL ISSUES

| | | Yes | No |
|---|---|---|---|
| **43.** | Do you need any help with your personal affairs? | ☐ | ☐ |
| **44.** | Would you like to talk to another professional about your condition or treatment? | ☐ | ☐ |
| **45.** | **Would you like any more information about the following?** | | |
| | a. Your condition | ☐ | ☐ |
| | b. Your care | ☐ | ☐ |
| | c. Your treatment | ☐ | ☐ |
| | d. Other types of support | ☐ | ☐ |
| | e. Financial issues | ☐ | ☐ |
| | f. Other (please state): ______________________ | | |

**Are there any other concerns that you would like us to know about?**

*Carry on over the page if needed*

**You can use this section to jot down any questions that you want to ask your doctors or other caring professionals**

Question 1

Question 2

Question 3
