## Supplementary material for "Patient reported distress can aid clinical decision making in idiopathic pulmonary fibrosis: analysis of the PROFILE cohort"

**Supplementary Document 2**

**Building IRT models: supplementary methods**

For SPARC questions that aimed to provide a metric of distress, two-parameter Item Response Theory (2PL-IRT) models were constructed for polytomous responses: 'Not at all', 'A little bit', 'Quite a bit', 'Very much' coded as 0, 1, 2, and 3 respectively, suggestive of monotonicity. For each model, exploratory factor analysis verified items as unidimensional and locally independent with one dominant factor, where no dominant factor was confirmed. Further breakdown and assessment of items was performed until suitable models were built. Model fit of data according to partial credit, graded response or rating scale was confirmed with Akaike's and Bayesian information criterion derived from likelihood-ratio tests.

Category characteristic curves and boundary characteristic curves were used to visually select items (survey questions) that showed the greatest discrimination (*a*) between grades of response, improving the ability to identify associations with the latent trait (θ). Although less intuitive for our purposes, item difficulty (*b*) refers to the *theta* value, where the probability of a response is 0.5. The selected items were used within the model to derive an individual's *theta* value for the latent trait e.g. distress from physical symptoms; a greater value indicates more distress.

Identifying differential item functioning is an essential aspect of IRT model building to remove demographic bias that is independent from the clinical context being assessed. An ordinal logistic regression strategy was used to detect differential item functioning, assessing uniform and non-uniform differential item functioning and addressing invariance. We assessed whether gender bias was responsible for differences in response despite the same underlying level of distress. We removed any items from our models that showed differential item functioning according to gender; ethnicity was not assessed as the PROFILE cohort is predominantly white-british.

To confirm the unidimensional assumption for 41 polytomous items, the model was initially built at the level of the item domains. Factor analysis offered no clear dominant group when all 'physical' items were used in the model. Assessment of item characteristic curves identified seven items from a total of 21 showing good discrimination along the latent trait: 7 (shortness of breath), 13 (feeling weak), 14 (feeling tired), 16 (feeling sleepy during the day), 17 (loss of appetite), 21 (feeling restless and agitated), 22 (feeling that symptoms are not controlled). Additionally, a decision was made to include item 41 (side effects) in the physiological model, though originally classed in the treatment domain and showing poor low discriminatory value, as no other item offered insight into distress of physical side effects. A graded response model was subsequently fit using these eight items that represented one dominant factor accounting for 47% of variance, and no differential item functioning in model detected. 'Feeling tired' had the highest discrimination value (*a*) of 3.91 (95%CI 2.69-5.13), meaning item-level scores clearly discriminated along the latent scale; 'side effects' had a low *a* value of 0.57 (95%CI 0.15-0.98) (Supplementary Table 3).

Characteristic curves identified five items from the 'psychological' items showing good discrimination from a total of nine: 23 (feeling anxious), 24 (low mood), 25 (feeling confused), 26 (unable to concentrate), 28 (everything is effort). In addition, we included items in later domains that we considered related to psychological distress, 32 (thoughts of death), originally classed in the spiritual domain; item 38 (worry about effects on family) originally classed in the social domain; and 42 (worry about long term effects) originally classed in the treatment domain. Item selection resulted in one dominant factor accounting for 46% of variance, confirming unidimensional items. A graded response model was fit using these eight items, with no differential item functioning detected in model.

Characteristic curves identified all 'independence' items as showing good discrimination and represented a single dominant factor accounting for 79% of variance: 33 (losing independence), 34 (ability to carry out daily activities), 35 (ability to carry out household tasks). Likelihood-ratio tests identified the rating scale model as fitting the independence items best, with no differential item functioning detected in model.

Two-tailed t-tests were used to identify any differences in the mean theta values derived from the models according to one year mortality (Supplementary Table 2). Where significance was detected, items were included in a final model, as this indicated differences in distress and SPARC response. Items from physical and independence domains were combined into a final model to measure the latent trait of distress from disease status in our cohort of people with IPF. No differential item functioning was detected in this selection, confirming invariance assumption. Factor analysis confirmed unidimensional items and addressed local independence, identifying a single dominant factor accounting for 44% of the variance. Likelihood-ratio tests identified the graded response model as the best fit for the overall data, with *a* values ranging from 0.56 to 3.12. Monotonicity was confirmed by probability of more extreme responses increasing across the scale of theta (Supplementary Figure 1).

**Supplementary Table 1: Item discrimination in models**

| Item | Description | *a* | 95%CI | *a* [1] | 95%CI |
|---|---|---|---|---|---|
| Latent ability i: physical symptom distress from | | | | | |
| 7 | Shortness of breath | *1.88* | 1.42-2.35 | *2.12* | 1.60-2.63 |
| 13 | Feeling weak | *3.32* | 2.40-4.24 | *3.12* | 2.29-3.95 |
| 14 | Feeling tired | *3.91* | 2.69-5.13 | *2.89* | *2.13-3.65* |
| 16 | Feeling sleepy in the day | *1.60* | 1.19-2.00 | *1.35* | 0.99-1.72 |
| 17 | Loss of appetite | *1.52* | 1.05-1.99 | *1.65* | 1.16-2.13 |
| 21 | Restless and agitated | *1.57* | 1.09-2.05 | *1.44* | 0.99-1.88 |
| 22 | Uncontrolled symptoms | *1.66* | 1.19-2.12 | *1.68* | 1.21-2.14 |
| 41 | Side effects of treatment | *0.57* | 0.15-0.98 | *0.56* | 0.16-0.97 |
| Latent ability ii: psychological symptom distress from | | | | | |
| 23 | Feeling anxious | *2.51* | 1.80-3.23 | *(excluded)* | |
| 24 | Low mood | *3.25* | 2.21-4.29 | *(excluded)* | |
| 25 | Feeling confused | *1.83* | 1.17-2.50 | *(excluded)* | |
| 26 | Unable to concentrate | *2.13* | 1.45-2.81 | *(excluded)* | |
| 28 | Everything is effort | *1.80* | 1.31-2.29 | *(excluded)* | |
| 32 | Thoughts of death and dying | *1.46* | 0.97-1.95 | *(excluded)* | |
| 38 | Worry about effect on family | *1.32* | 0.92-1.72 | *(excluded)* | |
| 42 | Worry about long term effects of treatment | *0.98* | 0.57-1.38 | *(excluded)* | |
| Latent ability iii: independence distress from | | | | | |
| 34 | Losing independence | | | *1.57* | 1.12-2.03 |
| 35 | Ability to carry out daily activities | *2.89* | 2.25-3.54 | *1.72* | 1.20-2.25 |
| 36 | Ability to carry out household tasks | | | *1.87* | 1.34-2.39 |

Item numbers included in IRT models presented, a: discrimination of item, [1]: discrimination in overall model. 95%CI: 95 % confidence interval for coefficient of a. Latent trait i-ii tested in graded response model, iii tested in rating scale model. Overall trait tested in graded response model.

**Supplementary Table 2: Survival bias in people with complete lung function recording**

| Lung measure | | FVC | | DLCO | |
|---|---|---|---|---|---|
| | *All* | *Missing* | *Complete* | *Missing* | *Complete* |
| Total | 243 | 21 | 222 | 75 | 168 |
| % Died | 20.2 | 61.9 | 16.2 | 34.7 | 13.7 |
| $\chi^2$ p value | | <0.0001 | | <0.0001 | |
| Mean distress | 0.0 | 0.17 | -0.02 | 0.28 | -0.13 |
| ttest p value | | 0.384 | | 0.002 | |
| Exp. Score (11Q:33) | 7.7 | 8.8 | 7.6 | 9.5 | 7.0 |

Chisquared ($\chi^2$) p-values based on observed and expected proportions alive and died between those missing lung function and those with complete lung function measures. Two-way ttest p value based on difference between mean theta values of those missing lung function and those with complete lung function measures. Exp. Score: expected score based on number of questions (nQ) and max score where 0=Not at all, 1=A little bit, 2=Quite a bit, 3=Very much.

**Supplementary Table 3: Difference in distress models according to one year mortality**

| **Mortality in 1 year:** | **Alive** | **Died** | ***(p-value)*** |
|---|---|---|---|
| Number | 194 | 49 | |
| Latent ability: physical symptom distress | | | |
| Mean Theta (θ) | -0.08 | 0.32 | ***0.0072*** |
| Std. Dev. | 0.92 | 0.92 | |
| Exp. Score (8Q:24) | 6.05 | 7.93 | |
| Latent ability: psychological symptom distress | | | |
| Mean Theta (θ) | -0.03 | 0.13 | *0.25* |
| Std. Dev. | 0.91 | 0.83 | |
| Exp. Score (8Q:24) | 3.33 | 3.93 | |
| Latent ability: independence distress | | | |
| Mean Theta (θ) | -0.07 | 0.33 | ***0.0024*** |
| Std. Dev. | 0.80 | 0.94 | |
| Exp. Score (3Q:9) | 0.62 | 1.49 | |
| Latent ability: overall distress | | | |
| Mean Theta (θ) | -0.10 | 0.39 | ***0.0012*** |
| Std. Dev. | 0.92 | 0.92 | |
| Exp. Score (11Q:33) | 7.2 | 10.2 | |

Two-tailed t-test p-values based on difference in mean theta values. Significant p-values ($p<0.05$) in bold. Std. Dev.: standard deviation from mean theta value. Exp. Score: expected score based on number of questions (nQ) and max score where 0=Not at all, 1=A little bit, 2=Quite a bit, 3=Very much.

**Supplementary Figure 1: Item characteristic curves of final model**

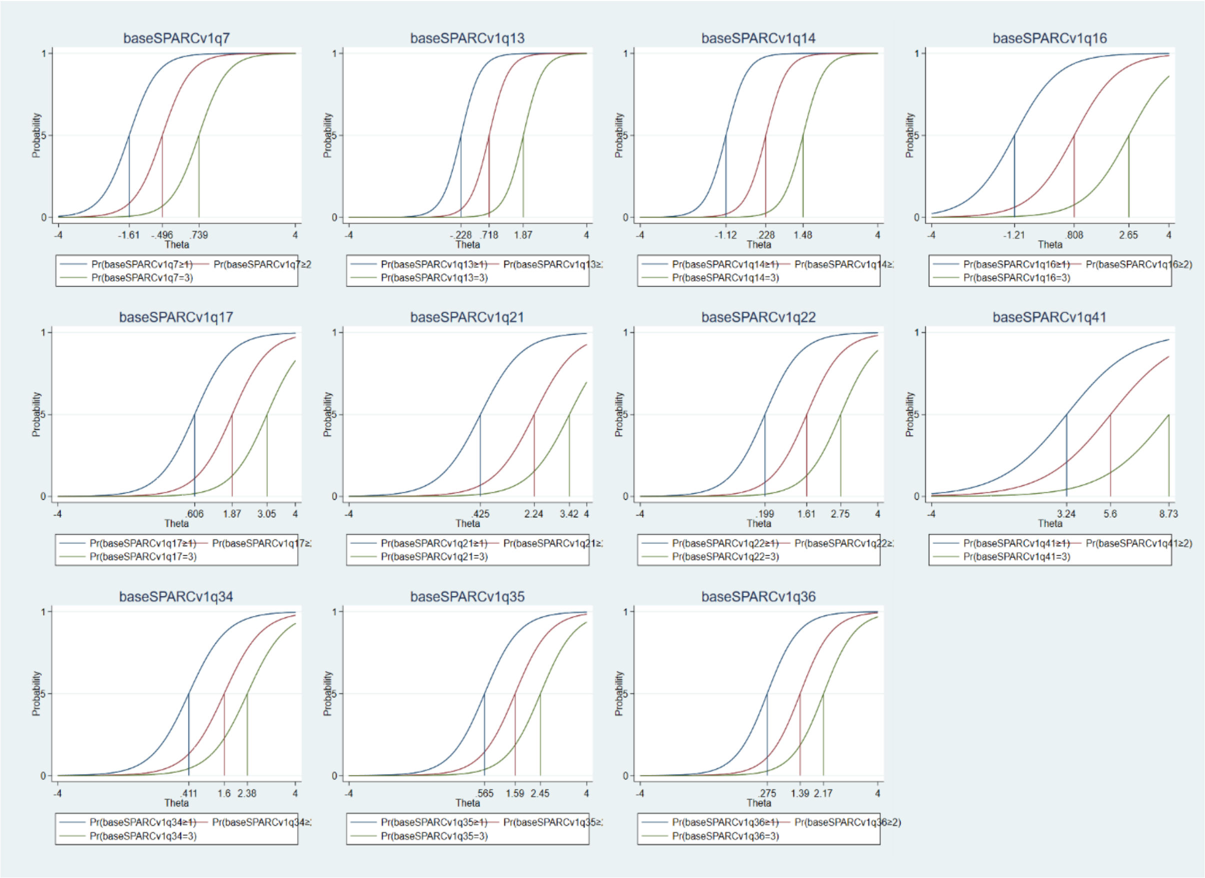


**Supplementary Figure 1 legend**: Item characteristic curves confirm monotonicity, 50% probability of a more extreme response increases along the latent trait. Good item discrimination is also evidenced by steep curves and little overlap.

**Supplementary Figure 3. ROC analysis of I-PARC ability to predict death where observations with missing lung function were included.**

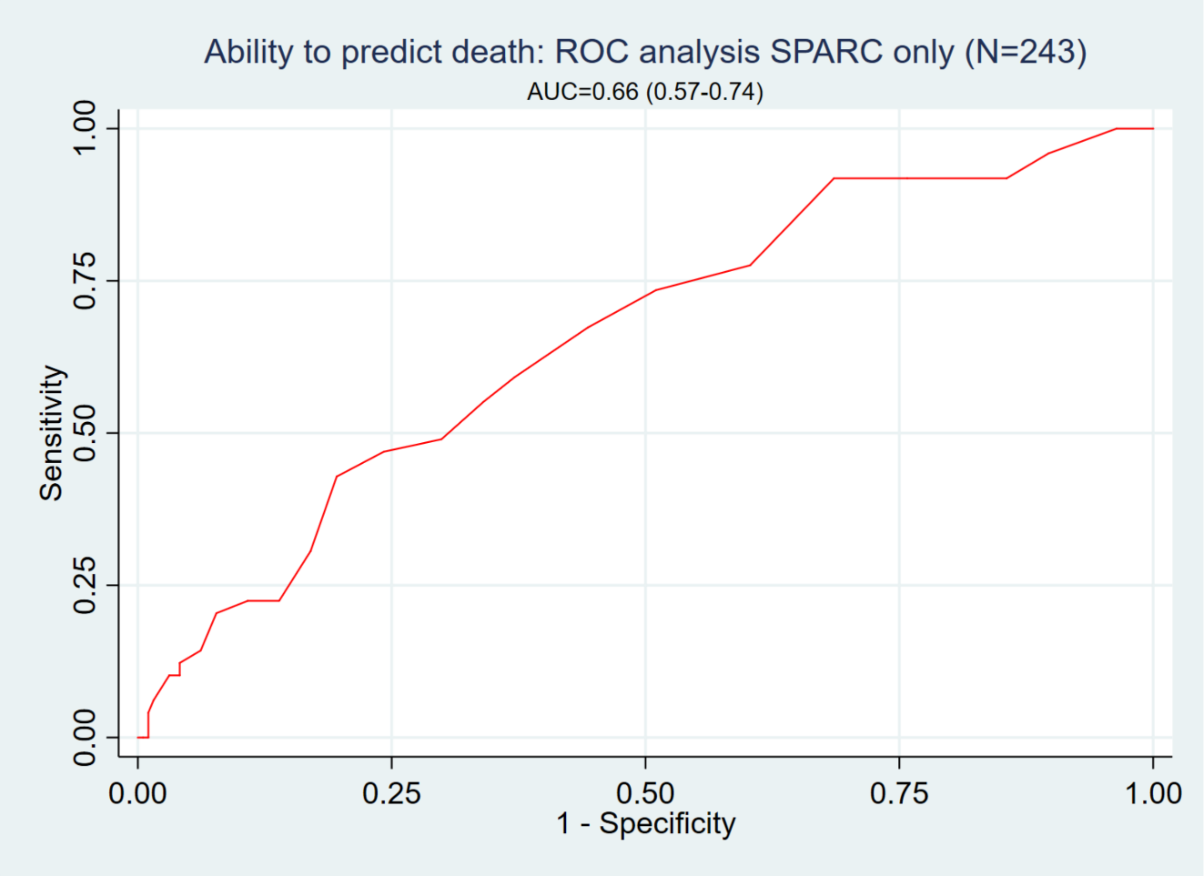


**Supplementary Figure 2 legend:** Area Under Curve (AUC) for the ability to predict one year mortality in 243 patients with available distress score was 0.66 (95%CI 0.57-0.74).

**Supplementary Figure 3: IPARC modification following pirfenidone intervention**

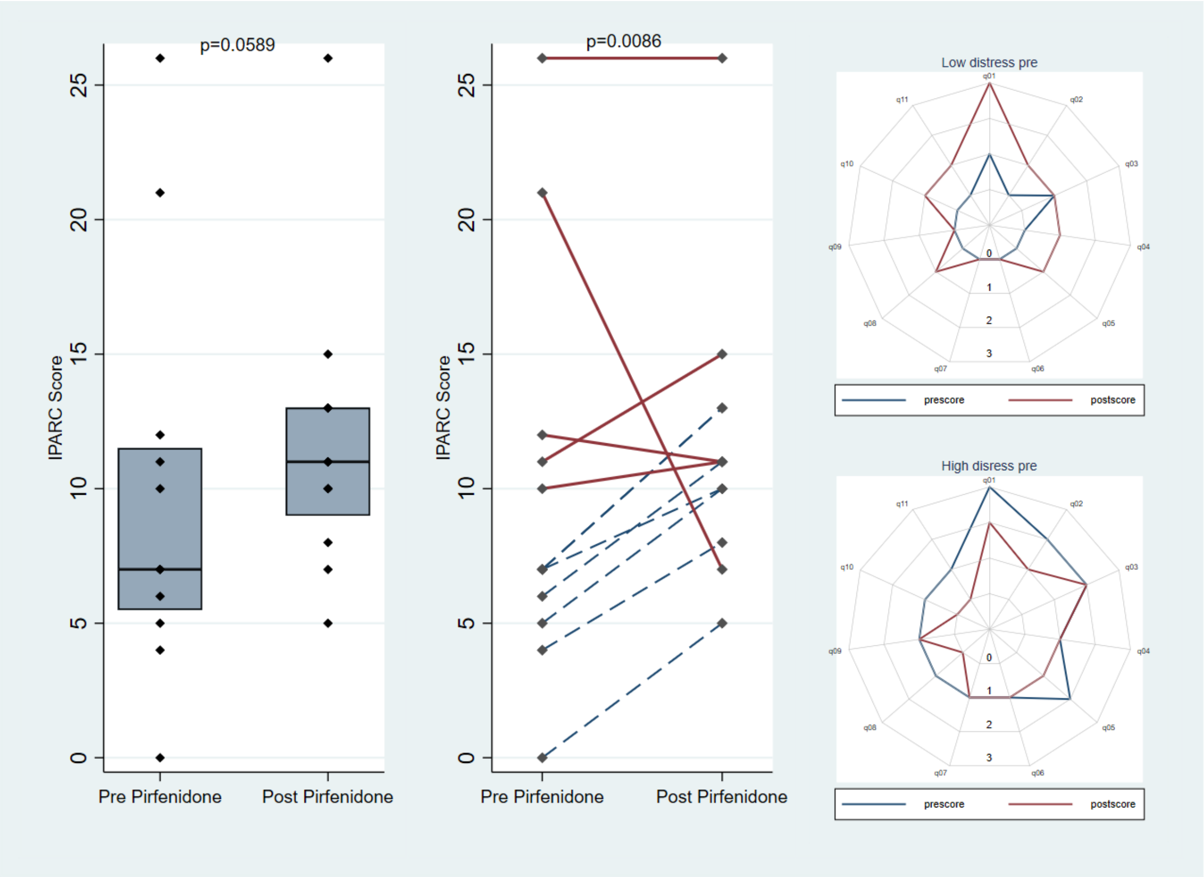


**Supplementary Figure 3 legend**: Overall median IPARC scores (IQR) for patients pre and post pirfenidone intervention, Wilcoxon signed-rank test indicates that IPARC scores following intervention were not significantly different (p=0.0589). The difference between post and pre intervention scores were associated with initial distress level (red solid lines - above threshold of 9; blue dashed line – below threshold of 9) (Wilcoxon rank sum p=0.0086). Radar plots of median pre and post scores according to initial distress level indicate how items are reported differently.
